# Prediction of Deleterious Single Nucleotide Polymorphisms in Human p53 Gene

**DOI:** 10.1101/408476

**Authors:** Saruar Alam, Mohammad Sayem, Md. Kamrul Hasan, Zinat Sharmin, Mahmud Arif Pavel, Md. Faruk Hossain

## Abstract

With a variety of accessible Single Nucleotide Polymorphisms (SNPs) data on human p53 gene, this investigation is intended to deal with detrimental SNPs in p53 gene by executing diverse valid computational tools, including Filter, SIFT, PredictSNP, Fathmm, UTRScan, ConSurf, Phyre, Tm-Adjust, I-Mutant, Task Seek after practical and basic appraisal, dissolvable openness, atomic progression, and analysing the energy minimization. Of 581 p53 SNPs, 420 SNPs are found to be missense or non-synonymous and 435 SNPs are in the 3 prime UTR and 112 SNPs are of every 5 prime UTR from which 16 non synonymous SNPs (nsSNPs) as non-tolerable while PredictSNP package predicted 14 (taking consideration SNP colored green by two or more than 2 analyses is neutral). By concentrating on six bioinformatics tools of various dimensions a combined output is generated where 14 nsSNPs are prone to exert a deleterious effect. By using diverse SNP analysing tools we have found 5 missense SNPs in the 3 crucial amino acids position in the DNA binding domain. The underlying discoveries are fortified by I-Mutant and Project HOPE. The ExPASy-PROSITE tools characterized whether the mutations located in the functional part of the protein or not. This study provides a decisive outcome concluding the accessible SNPs information by recognizing the five harming nsSNPs: rs28934573 (S241F), rs11540652 (R248Q), rs121913342 (R248W), rs121913343 (R273C) and rs28934576 (R273H). The findings of this investigation recognize the detrimental nsSNPs which enhance the danger of various kinds of oncogenesis in patients of different populations’ in genome-wide studies (GWS).

## 1. Introduction

Single Nucleotide Polymorphism (SNP) is the most prevalent form of genetic mutation occurs in human. About 93% of human genes are represented by single SNP [1]. SNPs are responsible for originating most of the variations among the people. SNPs can be in the coding regions, non-coding regions, and in the intergenic area between two genes [2]-[3]. Although coding SNPs and non-coding are phenotypically neutral, nsSNPs are supposed to influence phenotype by altering protein sequence [1]-[4]. As nsSNPs alter the amino acids in their corresponding protein, it could have deleterious effects on the structure and function [5]-[6]. They are associated with various human diseases and disorders. Several studies confirm the association of nsSNPs in susceptibility to infection and the progression of autoimmune diseases and inflammatory disorders [5]-[9]. About 50% of the mutations inclined with hereditary genetic disorders are nsSNPs [11]-[13]. As a result, many of the researchers are focusing on nsSNPs in cancer biology, precisely, in cancer-causing genes.

Mutations in the tumor suppressor gene p53 account for ∼50% of human cancers [17]-[18]. p53 is a critical regulator of tissue homeostasis [19] which binds to stabilize DNA as a tetramer that leads to the regulation of genes mediating key cellular processes including cell-cycle arrest, DNA repair, senescence and apoptosis [15]-[16]. The regular allele of p53 encodes a 53-kD nuclear phosphoprotein which plays an important role controlling cell proliferation [20]-[23]. However, in human tumors, point mutations, rearrangements, allelic loss and deletions are found in the p53 gene [24]-[27]. These abnormalities, together with the changes of oncogenes and tumor suppressor genes, consist the network of mutations that leads to malignancy. Despite the importance of p53, no computational studies have been reported that detects the deleterious nsSNPs in p53 gene. Therefore, in this current investigation, we carried out an in-silico analysis of p53 gene in order to characterize the deleterious mutations. Our study encompasses (i) retrieving SNPs in p53 gene from available databases, (ii) Allocating deleterious nsSNPs to their phenotypic effects based on the sequence and structure-based homology and identifying the regulatory nsSNPs responsible for altering the patterns of splicing and gene expression, (iii) predicting the role of the amino acids substitution on the secondary structures based on solvent stability and accessibility, and (iv) predicting the effect of mutations in the domain structures.

## 2. Materials and Methods

The Flow Chart expounds the whole process of our study in identification and characterization of detrimental SNPs in p53 gene. Both the structural and functional consequences have been analysed upon missense mutation (Figure 1).

**Figure 1:**
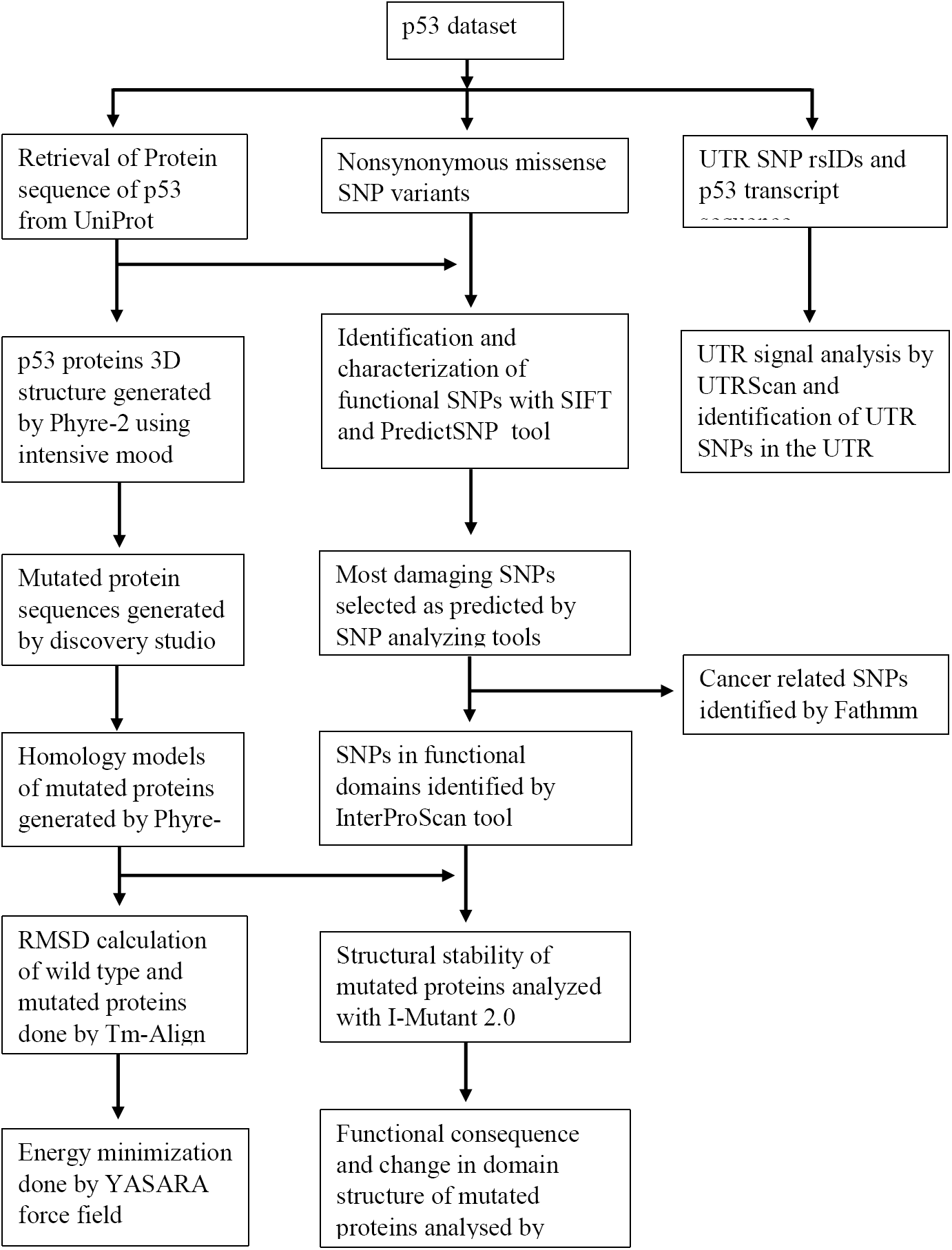
Flow chart of method and materials.

### 2.1. Retrieval of SNP Datasets

Data for human p53 gene was retrieved from an online web-based data sources such as Online Mendelian Inheritance in Man (OMIM) [28]. The SNPs information both protein accession number and SNP ID of the p53 gene was retrieved from the NCBI dbSNP (Database of Single Nucleotide Polymorphism) [29], and the protein fasta sequence was retrieved from UniProt (http://www.uniprot.org/) [30]

### 2.2. Analysis of Functional Consequences of nsSNPs

To sort out which SNP is Intolerant and which one is tolerant an online tool SIFT was employed to identify the deleterious coding nonsynonymous SNPs [31]. This program assumes that major amino acids will be conserved in the protein family and changes at particular positions have a tendency to be anticipated as harmful [31]-[32]. During mutagenesis studies in human SIFT can easily differentiate between functionally neutral or innocuous and detrimental polymorphisms [33]. SIFT programs algorithm was designed to use SWISSPROT, nr, and TrEMBL databases to find out homologous sequences with a query. In our study, the rsIDs (identification number) of each SNP of the human p53 gene obtained from NCBI were submitted to SIFT as a query sequence for homology searching. The SIFT score ≤0.05 was set to indicate the deleterious effect of a nonsynonymous mutation on protein function.

### 2.3. Characterization of Functional nsSNPs

For characterization of functional nsSNPs, we used PedictSNP web server (http://loschmidt.chemi.muni.cz/predictsnp) [34]. It was constructed from 3 independent datasets by eliminating all inconsistencies, duplicities, and mutations that were used before in the preparation of assessed tools. The standard dataset comprising over 43,000 mutations was taken for the impartial assessment of eight well-known prediction tools: nsSNPAnalyzer, MAPP, PANTHER, PolyPhen-1, PhD-SNP, PolyPhen-2, SIFT, and SNAP. The 6 best-performing tools were shared into an accord classifier PredictSNP, ensuing into drastically better prediction implementation, and simultaneous time returned results for all mutations, approving that unanimity prediction denotes an accurate and vigorous alternate to the predictions delivered by individual tools [34].

### 2.4. Prediction of Cancer Promoting Mutations

Some mutation may have an association with cancer. To predict cancer-associated SNP of p53 gene, we used the Functional Analysis through Hidden Markov Models (Fathmm) web server available at (http://fathmm.biocompute.org.uk/cancer.html) [35]. Fathmm allows to combine sequence conservation within hidden Markov models (HMMs) [36]. Fathmm server is well known as a high-throughput web server that is often employed to anticipate phenotypic, molecular, and functional consequences of protein variants on coding as well as the noncoding region. Fathmm employs two algorithms unweighted, sequence/conservation based and weighted, combined by sequence conservation with pathogenicity weights. In Fathmm server the default prediction threshold is set at −0.75 where prediction with a score less than this value predicts that the mutation is considered to be potentially associated with cancer. Cancer-promoting mutations is detrimental to our body, these types of mutations play a critical role in cell cycle regulation and mutations falling in the conserved region can depress the nature of the domain.

### 2.5. Identification of Functional SNPs in Conserved Regions

Functional amino acids remain conserved throughout evolution. Evolutionarily conserved amino acid residues in the p53 protein were anticipated by ConSurf web server available at (http://consurf.tau.ac.il/2016/index_proteins.php) [37] by using a Bayesian algorithm (conservation scores: 1–4 variable, 5-6 intermediate, and 7–9 conserved) [38]-[39]. Protein Fasta sequence was submitted and the conserved regions were predicted and shown by means of coloring scheme and conservation score of the amino acids. It also predicts the functional and structural residues of the protein. Highly conserved amino acids located at high-risk nsSNP sites were selected for further analysis.

### 2.6. Scanning of UTR SNPs

UTRs are known to play vital roles in the post-transcriptional instruction and regulation of gene expression, comprising modulation of the transport of mRNAs out of the nucleus and of translation competence, subcellular localization, and constancy [40]. To find the functional SNPs we employed UTRScan a web server [41], an alignment matcher which searches nucleotide (RNA, tRNA, DNA) or protein sequences to find UTR motifs and is able to locate, in a particular sequence, motifs that distinguish 3’UTR and 5’UTR sequences. Such motifs are well-defined in the UTRSite Database, a compilation of functional sequence arrangements located in the 5’- or 3’-UTR sequences [42]-[43]. If a SNP with a different nucleotide at each UTR is found to have dissimilar working patterns, this UTR SNP is expected to have an impact on the mRNA stability. To perform this, 5’- and 3’-UTR SNPs from NCBI were submitted in FASTA format and the results showed predicted UTRs at the specific region.

### 2.7. Identification of a deleterious mutation in the functional domain

The functional domain of the product of p53 gene was identified using InterProScan. InterProScan connects diverse protein signature identification methods from the InterPro consortium associate databases into one resource. A web-based version of InterPro is accessible for academic and profitable organizations from the EBI (http://www.ebi.ac.uk/InterProScan/) [44]. Using InterProScan tool allows scanning protein sequence for matches against the InterPro protein signature databases. InterProScan takes protein sequence in FASTA format. After analyzing the deleterious mutation from the SIFT mutation among them which mutation is in the functional domain of the p53 protein is identified.

### 2.8. Modelling of the Mutated Protein

Phyre2 (Protein Homology/ Analogy Recognition Engine) [45], one of the most common online protein fold detection server, was employed to generate a 3D model of the protein. Virtual Mutation (VM) denotes the substitution of a single or multiple amino acids in the atomic 3D model of the molecule [46]-[47]. Accelrys Discovery Studio 4.0 was employed to create a mutated sequence for the corresponding amino acid change [48]. Regenerated mutant sequences were used further for the purpose of mutant modelling. The modelling of mutated proteins was also performed through, Phyre2 Protein Fold Recognition Server. Phyre2 nominate the best-matched template and generate a model through successive phases, such as similarity analysis, profile construction, and structural properties. We selected intensive mode of protein modelling to obtain an accurate model. The input sequence of the proteins was given in FASTA layout.

### 2.9. Energy Minimization and RMSD Calculation of the Protein Models

TM-align is an algorithm used for the protein structure similarities. TM-align combines the TM-score rotation matrix and Dynamic Programming (DP) to identify the best structural alignment between protein pairs. This server was used for RMSD calculation of the protein structures [49]. YASARA-minimization server was employed to perform the energy minimization of the generated mutated protein models. YASARA (Yet Another Scientific Artificial Reality Application) is a modelling and simulation software with variety of application. YASARA-minimization server uses YASARA force field for energy minimization that can optimize the damage of the mutant proteins and thus precisely calculates the energy that is reliable. To perform this task, the PDB file of the mutant proteins were inserted as input data and the result was additionally examined for imminent advance [50].

### 2.10. Effect of Mutation in Protein Stability

I-Mutant 3.0 server was employed to perform the prediction of the alteration in stability upon mutations. I-Mutant is a high throughput support vector machine (SVM) based tool server, which can automatically anticipate the alteration in stability of the structure by examining the structure of the protein sequence. I-Mutant 3.0 can be utilized as a classifier to predict the sign of stability with mutations and a regression calculator which anticipates the distinction in Gibbs free energy. The resulting DDG value indicates the difference between the Gibbs free energy of mutated protein and wild-type protein in kcal/mol.[51].

### 2.11. Prediction of Structural Effects upon Mutation

Project HOPE (Project Have your Protein Explained) was employed to understand the effect of the amino acid substitutions [52]. HOPE (http://www.cmbi.ru.nl/hope/input/) server was utilized for molecular dynamics simulation to observe the effect of the mutations on the structure of p53. This web server performs a BLAST against the PDB and builds a homology model of the query protein if possible through YASARA and collects 3D structure data from What IF web services and then get the sequence from the UniProt database and get features like an active site, motifs, domains and so forth. Finally, to predict the features of the protein it uses Distributed Annotation System (DAS) servers through which it can exchange annotations on genomic and protein sequences [53].

## 3. Results and discussions

### 3.1 SNP database from for p53

The polymorphism data is available in many databases, NCBI dbSNP housed extensive data for different gene NCBI has the largest database that is helpful for analysing single nucleotide polymorphism. 581 SNPs have been found for cellular tumor antigen p53, (NCBI Reference Sequence: NP_000537.3) among them 420 SNP found to be missense or nonsynonymous among 581 there are 435 SNP in 3 prime UTR and 112 in 5 prime UTR region. Only the missense, 3 prime, and 5 prime UTR SNPs have been selected for further analysis.

### 3.2 Prediction of detrimental nonsynonymous SNP

An online tool SIFT was employed which analyse conservancy of an amino acid by sequence homology to determine the conservation of a specific position of any amino acids in a protein. SIFT does the alignment between paralogous and orthologous protein’s amino acid sequences to determine the effect of an amino acid replacement analysing its functional importance and physical characteristics. SIFT takes rsID as input. In our analysis, we found 16 missense SNPs among 420 to be predicted as deleterious by SIFT webserver.

### 3.3 Analysis of SIFT predicted deleterious SNPs

In SIFT analysis, 16 SNPs have been found to be detrimental. These 16 SNPs was also analyzed in an online SNP analyzing tool PredictSNP which is a package system where other SNP analyzing method has been assembled. PredictSNP takes input protein sequence as FASTA format and then the SIFT result mutation was done in PredictSNP and the impact of the mutation of analyzed by choosing nsSNPAnalyzer, MAPP, PANTHER, PolyPhen-1, PhD-SNP, PolyPhen-2, SNAP and SIFT tool. PredictSNP shows result of different SNP in which percentage they are detrimental or neutral the percentage value indicates the confidence of the result. SNPs that were found to be predicted as neutral by more than one tool have been excluded from our study.

### 3.4 Identification of cancer associated SNPs from predicted deleterious SNPs

SNPs that were predicted as deleterious were then analysed using Fathmm to test whether they are associated with cancer or not. In our analysis, we have found that each and every SNPs that were predicted as detrimental are also predicted as having their association with cancer. In Fathmm server, the default prediction threshold is –0.75 where prediction with a score less than this indicates that the mutation is potentially associated with cancer. In our result, we find that predicted score is very low than the threshold that indicates that there is much higher potentiality of having a relation of these SNP in cancer.

### 3.5 Identification of functionally important SNPs in the conserved regions

Some amino acids are found to be crucial for the function of a protein. Essential amino acids for showing function are tend to be evolutionary conserved. To identify the evolutionarily conserved amino acids along the protein we used the online tool consurf. Consurf gives result in a tabulated form where conserved amino acids in a specific position are shown with CS and color value where the lower the CS and higher the color value indicates the higher conservancy.

### 3.6 Functional SNPs in UTR identification

3 prime UTR region has a significant effect on gene expression due to the defective ribosomal RNA translation or by affecting RNA half-life. 5 prime UTR also play important role in mRNA stabilization. The UTRscan server was used to analyze 77 3’UTR SNPs and 129 5’ UTR SNPs of p53 gene. The UTRscan server looks for patterns in the UTR database for regulatory region motifs and according to the given SNP information predicts if any matched regulatory region is damaged[54]. UTRscan found 8 UTRsite motif matches in the p53 transcript. Total 141 matches were found for 5 motifs

### 3.7 Prediction of deleterious mutation in the functional domain of p53

From InterproScan tool analysis we found that there are three functional domains in the p53 gene. InterProScan takes FASTA format of protein sequence as input and allows to scan matching protein sequence against the InterPro protein signature databases. InterProScan also predict the function of the residues. It also provides the consensus amino acid for protein function and whether the predicted deleterious mutation SNP is necessary or not for function. If the predicted deleterious SNP found to be in the amino acid in the functional domain or functioning residue then we can authenticate our prediction. The InterProScan result is tabulated in table 6.

**Table 1:** Prediction of deleterious SNP by SIFT tool

| SNP | Position | Amino acid change | Prediction | Score | Median |
| --- | --- | --- | --- | --- | --- |
| rs11540652 | 248 | R248Q | Damaging | 0.00 | 2.11 |
| rs11540654 | 110 | R110L | Damaging | 0.04 | 2.13 |
| rs11540654 | 110 | R110P | Damaging | 0.04 | 2.13 |
| rs17849781 | 278 | P278A | Damaging | 0.00 | 2.11 |
| rs28934571 | 249 | R249S | Damaging | 0.00 | 2.11 |
| rs28934573 | 241 | S241F | Damaging | 0.00 | 2.11 |
| rs28934574 | 282 | R282W | Damaging | 0.00 | 2.14 |
| rs28934576 | 273 | R273H | Damaging | 0.00 | 2.11 |
| rs28934577 | 257 | L257Q | Damaging | 0.00 | 2.11 |
| rs28934578 | 175 | R175H | Damaging | 0.00 | 2.11 |
| rs28934873 | 133 | M133T | Damaging | 0.00 | 2.13 |
| rs28934874 | 151 | P151T | Damaging | 0.02 | 2.11 |
| rs55832599 | 267 | R267W | Damaging | 0.00 | 2.11 |
| rs112431538 | 285 | E285K | Damaging | 0.01 | 2.14 |
| rs121913343 | 273 | R273C | Damaging | 0.00 | 2.11 |
| rs121913342 | 248 | R248W | Damaging | 0.00 | 2.11 |

**Table 2:** Further analysis of 16 SIFT predicted nsSNPs by PredictSNP online tool, predicted by different tools as deleterious are highlighted in the yellow mark and neutral in green color.

| SNP | Mutation | Predict SNP | MAPP | PhD-SNP | PolyPhen-1 | Polyphen-2 | SIFT | SNAP | nsSNP Analyzer | PANTHER |
| --- | --- | --- | --- | --- | --- | --- | --- | --- | --- | --- |
| rs11540654 | R110L | 76% | 72% | 82% | 59% | 47% | 46% | 50% | 63% | 48% |
| rs11540654 | R110P | 87% | 59% | 88% | 59% | 63% | 46% | 56% | 63% | 57% |
| rs28934873 | M133T | 51% | 77% | 61% | 67% | 76% | 79% | 50% | 63% | 56% |
| rs28934874 | P151T | 87% | 48% | 88% | 59% | 55% | 79% | 72% | 63% | - |
| rs28934578 | R175H | 87% | 63% | 88% | 59% | 56% | 79% | 72% | 63% | - |
| rs28934573 | S241F | 87% | 77% | 88% | 74% | 81% | 79% | 89% | 63% | - |
| rs11540652 | R248Q | 87% | 59% | 88% | 59% | 81% | 53% | 81% | 65% | - |
| rs121913342 | R248W | 87% | 77% | 88% | 74% | 81% | 79% | 85% | 65% | - |
| rs28934571 | R249S | 87% | 77% | 88% | 74% | 81% | 53% | 81% | 63% | - |
| rs28934577 | L257Q | 87% | 78% | 88% | 74% | 81% | 79% | 72% | 63% | - |
| rs55832599 | R267W | 87% | 77% | 88% | 74% | 65% | 79% | 81% | 63% | - |
| rs121913343 | R273C | 87% | 91% | 88% | 74% | 81% | 79% | 85% | 63% | - |
| rs28934576 | R273H | 87% | 77% | 88% | 59% | 65% | 53% | 62% | 65% | - |
| rs17849781 | P278A | 87% | 77% | 88% | 59% | 81% | 53% | 81% | 63% | - |
| rs28934574 | R282W | 87% | 77% | 88% | 74% | 81% | 79% | 72% | 63% | - |
| rs112431538 | E285K | 87% | 84% | 77% | 74% | 68% | 79% | 62% | 65% | - |

**Table 3:**
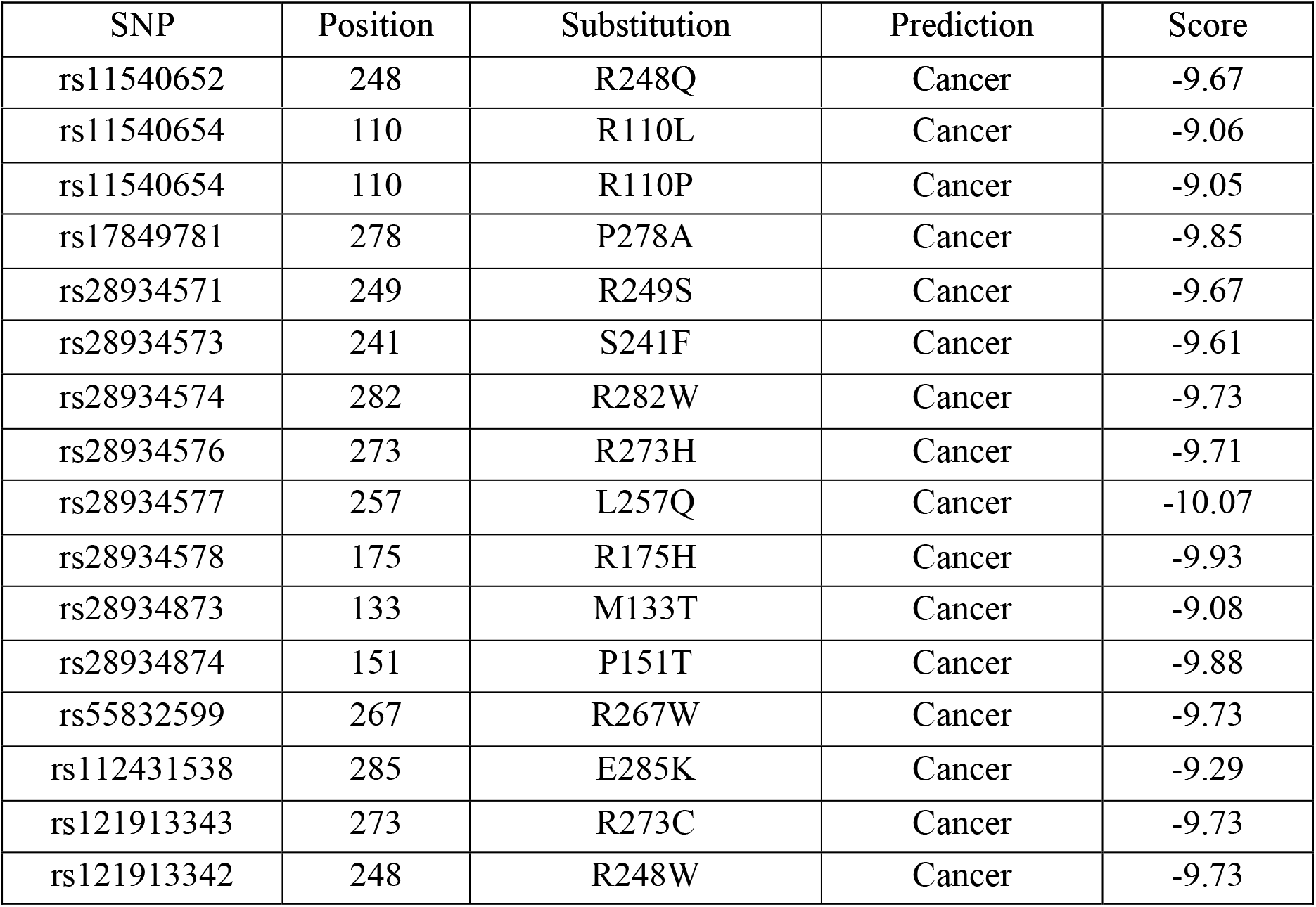
Cancer-associated SNP where the threshold value is −0.75. Scores having −0.75 predicted as having their association with cancer.

| SNP | Position | Substitution | Prediction | Score |
| --- | --- | --- | --- | --- |
| rs11540652 | 248 | R248Q | Cancer | -9.67 |
| rs11540654 | 110 | R110L | Cancer | -9.06 |
| rs11540654 | 110 | R110P | Cancer | -9.05 |
| rs17849781 | 278 | P278A | Cancer | -9.85 |
| rs28934571 | 249 | R249S | Cancer | -9.67 |
| rs28934573 | 241 | S241F | Cancer | -9.61 |
| rs28934574 | 282 | R282W | Cancer | -9.73 |
| rs28934576 | 273 | R273H | Cancer | -9.71 |
| rs28934577 | 257 | L257Q | Cancer | -10.07 |
| rs28934578 | 175 | R175H | Cancer | -9.93 |
| rs28934873 | 133 | M133T | Cancer | -9.08 |
| rs28934874 | 151 | P151T | Cancer | -9.88 |
| rs55832599 | 267 | R267W | Cancer | -9.73 |
| rs112431538 | 285 | E285K | Cancer | -9.29 |
| rs121913343 | 273 | R273C | Cancer | -9.73 |
| rs121913342 | 248 | R248W | Cancer | -9.73 |

**Table 4:** Summary of amino acids (conservation profile) in that corresponds in region with high-risk nsSNPs. CS: conservation score, color (6–9 = conserved, 5 = average, and 1–4 = variable); (s): predicted structural site; (f): predicted functional site.

| Residue position | Amino acid | CS | Color | B/E | function |
| --- | --- | --- | --- | --- | --- |
| 110 | R | 1.057 | 1 | E |  |
| 133 | M | -1.019 | 9 | B | S |
| 151 | P | -0.801 | 8 | E | F |
| 175 | R | -0.989 | 9 | E | F |
| 241 | S | -1.162 | 9 | E | F |
| 248 | R | -1.070 | 9 | E | F |
| 249 | R | -0.990 | 9 | E | F |
| 257 | L | -0.951 | 9 | B | S |
| 267 | R | -0.944 | 9 | E | F |
| 273 | R | -1.073 | 9 | E | F |
| 278 | P | -1.030 | 9 | E | F |
| 282 | R | -1.029 | 9 | E | F |
| 285 | E | -0.878 | 8 | E | F |

**Table 5:** Result showing UTR regions in the p53 transcript from UTRScan server.

| Signal name | UTR region | Match total | Position in transcript |
| --- | --- | --- | --- |
| uORF (Upstream open reading frames) | 5' | 44 | - |
|  | 3' | 25 | - |
| GY-BOX | 5' | 9 | - |
|  | 3' | 6 | - |
| IRES<br>(Internal ribosome entry site) | 5' | 17 | - |
|  | 3' | 11 | - |
| TOP<br>(Terminal oligopyrimidine) | 5' | 4 |  |
|  | 3' | 9 | - |
| UNR-bs | 3' | 7 | - |
| PAS<br>(Polyadenylation Signal) | 3' | 6 | - |
| K-BOX | 5' | 2 | 13-70<br>1-8 |
| ARE2 | 3' | 1 | 409-444 |

**Table 6:**
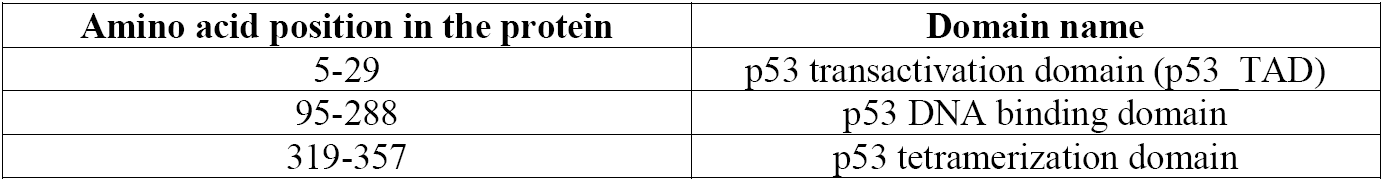
Functional domain and their position in p53 protein

### 3.8 Comparative Modeling of High-Risk Nonsynonymous SNPs and RMSD calculation

We used the Phyre2 protein model prediction tool to get the 3D structure human p53 protein with the predicted SNPs. Then introduced the following nonsynonymous mutations in the DNA binding domain: S241F, R248Q, R248W, R273H, R273C using Accelrys Discovery Studio 4.0. The models were then subjected to YASARA energy minimization server for energy minimization. Energy minimization results showed decreased free energy for all the mutant models than the wild-type models. The results are shown in Table 8. RMSD was calculated using the Tm-Align tool where the results were shown to be 2.99 for S241F, 1.97 for R273C and 1.45 for R248Q. These outcomes demonstrate a critical change in the proteins structure which can alter its natural function.

**Table 7:** Residue annotation of the protein

| Functioning site | Functioning residue with position |
| --- | --- |
| Zinc binding site | 176C, 179H, 238C, 242C |
| Dimerization site | 177P, 178H, 179H, 181R |
| DNA binding site | 239N, 241S, 248R, 273R, 275C, 276A, 277C, 280R |

**Table 8:** RMSD (Å) values and total free energy after energy minimization of the both wild type and mutated protein models

| Mutated models | Energy after minimization (kJ/mol) | RMSD (Å) |
| --- | --- | --- |
| Wild Type | -173964.6 |  |
| S241F | -155007.3 | 2.99 |
| R248Q | -165923.9 | 1.45 |
| R248W | -163338.0 | 1.45 |
| R273H | 5925809.8 | 1.97 |
| R273C | 7466100.3 | 1.97 |

After mutation of the wild-type, it was found that in every case energy after minimization is higher (more positive) than wild-type that indicates that these mutations destabilize the structure of the protein and in case of a mutation in the 273rd arginine amino acid position by histidine and cysteine affects the structure of the protein more.

### 3.9. Prediction of Protein Structural Stability

We used the neural network based routine tool I-Mutant 3.0 for studying the potential change in the stability of protein upon mutations. This tool took the input of the mutated protein models derived from PHYRE-2 server in PDB format. I-Mutant 3.0 creates results taking help of ProTherm database. This database housed the extensive amount of experimental data on free energy alterations due to mutations. In addition, this tool predicts score of free energy change due to mutations incorporating the energy based online tool FOLD-X. This increases the precision to 93% on one-third of the database if the FOLD-X analysis is incorporated along with I-Mutant[55]. Models with following mutations: S241F, R248Q, R248W, R273H, R273C were subjected to the server to predict DDG stability and RSA calculation. Result shows that everymutations decreased the stability of protein. Mutation R273H was responsible for the lowest DDG value (−1.62 kcal/mol) which followed by R273C (−1.52 kcal/mol).DDG values for other mutations ranged from −0.51 kcal/mol to −0.93 kcal/mol; this negative DDG values indicates decreased protein stability. The results are shown in Table 9.

**Table 9:**
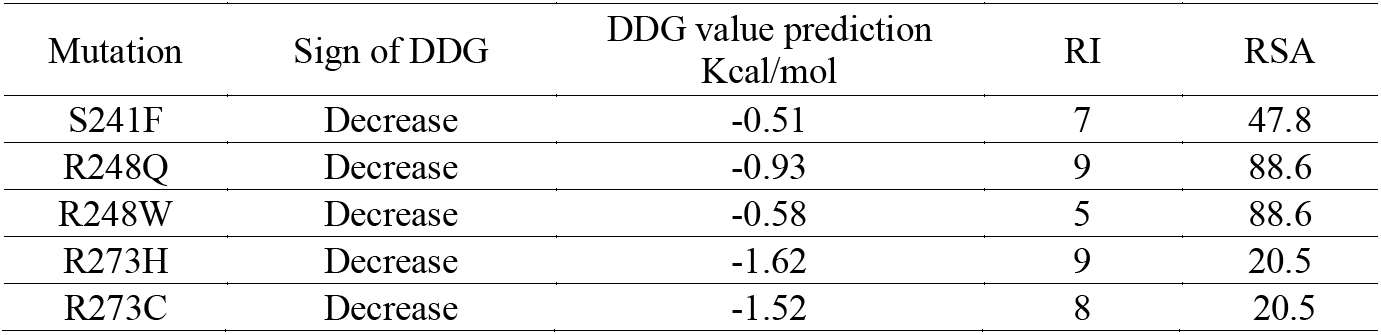
Table 7: I-mutant predictions for selected nsSNPs

| Mutation | Sign of DDG | DDG value prediction<br>Kcal/mol | RI | RSA |
| --- | --- | --- | --- | --- |
| S241F | Decrease | -0.51 | 7 | 47.8 |
| R248Q | Decrease | -0.93 | 9 | 88.6 |
| R248W | Decrease | -0.58 | 5 | 88.6 |
| R273H | Decrease | -1.62 | 9 | 20.5 |
| R273C | Decrease | -1.52 | 8 | 20.5 |

### 3.10 Analysis of structural effect upon mutation in DNA binding domain

The InterproScan tool was used to find the functional domain in p53 protein and map the predicted deleterious mutations in the functional domains for anticipating the changes they might cause in the domain structures. Among predicted 14 detrimental SNPs by different SNPs analyzer tool, we found that 5 missense SNP in the 3 important amino acids located in a domain which is responsible for DNA binding. These amino acids are essential for the functional activity of the domain. Therefore, a mutation in this amino acid position could change the protein structure as well as the function. We observe the effect on the structure of p53 under this 5 missense SNP using an online tool HOPE.

In figure 2A, the wild-type residue has positively charged Arginine (R). However the mutation from Arginine to Glutamine (Q) at 248^th^position makes the mutant is neutral. In figure 2B, Serine (S) mutated into Phenylalanine (F) at 241th position. The mutant residue is bigger and hydrophobic than the wild-type. In figure 2C, Arginine (R) mutated into Histidine (H) at 273th position and the mutant residue is smaller and neutral whereas wild-type is positively charged. In case of figure 2D, the Arginine(R) mutated into Cysteine (C) at 273th position and the mutant residue is smaller and neutral but the wild-type is positively charged. In the last figure 2E, Arginine (R) mutated into a Tryptophan (W) at 248^th^position. The mutant residue is bigger and neutral whereas wild type is bigger and positively charged.

**Figure 2:**
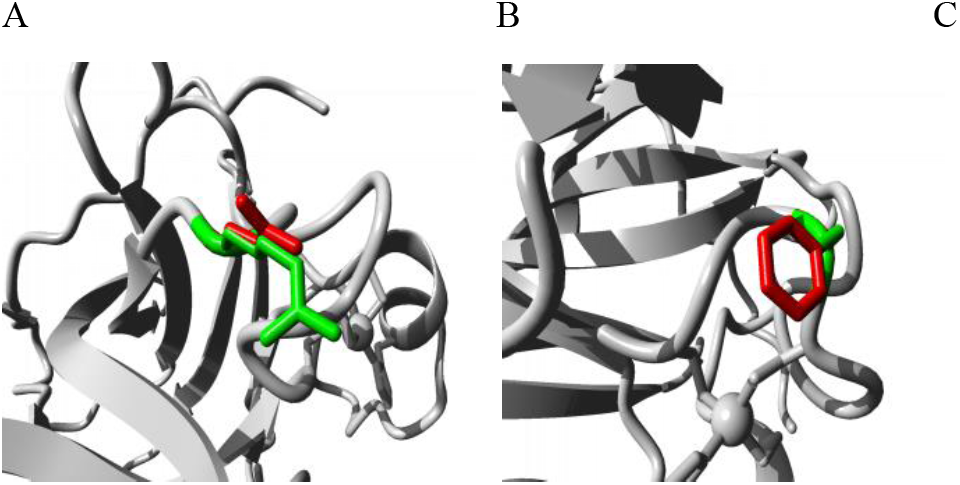
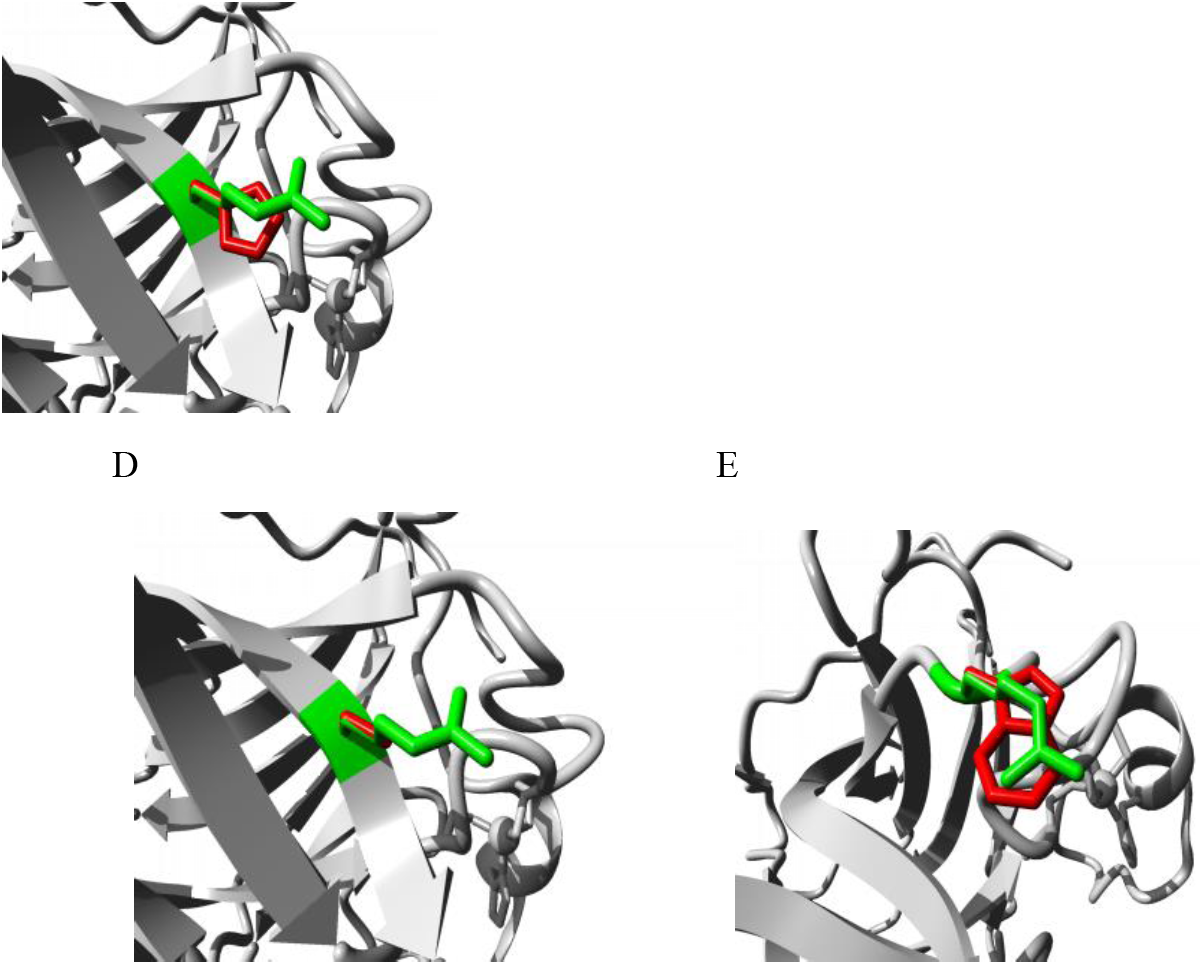
Close-up of the mutation A R248Q, B S241F, C R273H, D R273C, E 248W. (The protein = grey; side chains of wild type =green; mutant residue = red).

## Conclusion

In this study different SNP analysing tools has been employed for analysis of the available data from the NCBI dbSNP database for the tumor suppressor gene P53. The predicted deleterious SNPs were evaluated for their potential detrimental effects on the protein function and stability. Five SNPs were predicted deleterious; those are rs28934573 (S241F), rs11540652 (R248Q), rs121913342 (R248W), rs121913343 (R273C) and rs28934576 (R273H); and they have the most probability to increase P53 susceptibility. Henceforth, it is very likely that there are unreported nsSNPs increasing disease predisposition by altering protein function or structure. The findings of this study might be helpful in early diagnosing the detrimental SNPs that has probability to increase the risk of different types of cancer formation. Individuals diagnosed with the above nsSNPs can take precautions to avoid other risk factors associated with cancer establishment as they are susceptible to cancer due to these nsSNPs in P53; a major tumor suppressor gene. However, population based studies and wet lab experiments are beyond our scopes for the verification of the findings of the current study. Therefore, extensive and clinical studies are required to characterize the vast SNP data.

## Acknowledgment

The authors are grateful to the Department of Biochemistry and Molecular Biology, University of Dhaka, Bangladesh and the Department of Biological Sciences, St. John’s University, Queens, New York 11439.

